# Sustained responses and neural synchronization to amplitude and frequency modulation in sound change with age

**DOI:** 10.1101/2022.04.28.489911

**Authors:** Björn Herrmann, Burkhard Maess, Ingrid S. Johnsrude

## Abstract

Perception of speech requires sensitivity to features, such as amplitude and frequency modulations, that are often temporally regular. Previous work suggests age-related changes in neural responses to temporally regular features, but little work has focused on age differences for different types of modulations. We recorded magnetoencephalography in younger (21–33 years) and older adults (53–73 years) to investigate age differences in neural responses to slow (2-6 Hz sinusoidal and non-sinusoidal) modulations in amplitude, frequency, or combined amplitude and frequency. Audiometric pure-tone average thresholds were elevated in older compared to younger adults, indicating subclinical hearing impairment in the recruited older-adult sample. Neural responses to sound onset (independent of temporal modulations) were increased in magnitude in older compared to younger adults, suggesting hyperresponsivity and a loss of inhibition in the aged auditory system. Analyses of neural activity to modulations revealed greater neural synchronization with amplitude, frequency, and combined amplitude-frequency modulations for older compared to younger adults. This potentiated response generalized across different degrees of temporal regularity (sinusoidal and non-sinusoidal), although neural synchronization was generally lower for non-sinusoidal modulation. Despite greater synchronization, sustained neural activity was reduced in older compared to younger adults for sounds modulated both sinusoidally and non-sinusoidally in frequency. Our results suggest age differences in the sensitivity of the auditory system to features present in speech and other natural sounds.

## Introduction

Many middle-aged or older adults live with some form of hearing impairment (Cruickshanks et al., 1998; Feder et al., 2015; Goman and Lin, 2016) and often experience difficulties understanding speech in the presence of background sound (Pichora-Fuller, 2003; Pichora-Fuller et al., 2016). Speech contains regular and quasi-regular low-frequency (<10 Hz) fluctuations in amplitude and frequency (Rosen, 1992; Varnet et al., 2017). Sensitivity to such amplitude and frequency fluctuations is thought to be important for segregating concurrent sound as separate streams (Schröger, 2005, 2007; Snyder and Alain, 2007; Winkler et al., 2009; Bendixen, 2014) and for recognizing and predicting features of the acoustic environment (Jones and Boltz, 1989; Barnes and Jones, 2000; Jones et al., 2002; Henry and Herrmann, 2014; Nobre and van Ede, 2018). The current study is concerned with how low-frequency amplitude and frequency modulations are processed in the brain and whether processing of amplitude and frequency modulations differ between younger and older adults.

Sensitivity of the auditory system to temporally regular structure in sounds has been examined by focusing on at least two neural signatures: neural synchronization and sustained neural activity. Neural activity recorded using electro- or magnetoencephalography (EEG/MEG) synchronizes with regular amplitude and frequency modulations in sounds (Maiste and Picton, 1989; John et al., 2001; Boettcher et al., 2002; Picton et al., 2003; Henry and Obleser, 2012; Henry et al., 2014; Goossens et al., 2016; Herrmann and Johnsrude, 2018; Goossens et al., 2019; Herrmann et al., 2019; Bauer et al., 2021) and this neural synchronization increases with increasing coherence of a sound’s temporal modulation (Herrmann and Johnsrude, 2018). Synchronization of neural activity with regular structure in sounds is thought to provide the means by which neural circuits support generating predictions about the timing and features of future sounds (Henry and Herrmann, 2014) and filter out irrelevant sound information with different temporal dynamics (Lakatos et al., 2008; Lakatos et al., 2013; O’Connell et al., 2014; Obleser and Kayser, 2019).

The second neural signature of temporal regularity processing is a sustained response in EEG/MEG recordings (Barascud et al., 2016; Southwell et al., 2017; Herrmann et al., 2021). The sustained response is an ultralow-frequency response (i.e., a constant or slowly drifting offset) elicited by the occurrence of a temporally regular pattern in a sound. A sustained response has been demonstrated in response to repeated sets of pure tones in longer sequences of tones (Barascud et al., 2016; Southwell et al., 2017; Herrmann and Johnsrude, 2018; Southwell and Chait, 2018; Herrmann et al., 2022), coherent frequency changes in multitone chords (Teki et al., 2016; Dheerendra et al., 2021), and complex sounds made of isochronous tone sequences (Sohoglu and Chait, 2016). A sustained response has also been observed for amplitude (Ross et al., 2002; Herrmann et al., 2019) and frequency modulations in sounds (Herrmann and Johnsrude, 2018), and for isochronously presented clicks or vowels compared to irregular presentations (Gutschalk et al., 2002; Gutschalk and Uppenkamp, 2011). A sustained response elicited by temporally regular patterns in sounds has been suggested as a possible indicator of prediction-related processes (Heilbron and Chait, 2018).

Aging and hearing loss are associated with a loss of inhibition and an increase in excitation in the auditory system (Caspary et al., 1995; Caspary et al., 2008; Takesian et al., 2012; Knipper et al., 2013; Auerbach et al., 2014; Ouellet and de Villers-Sidani, 2014; Resnik and Polley, 2017; Salvi et al., 2017; Herrmann and Butler, 2021b) that are thought to result from damage to the auditory periphery and the associated deprived input to the auditory neural pathway (Zhao et al., 2016; Herrmann and Butler, 2021a). The consequence of reduced inhibition and increased excitation is hyperactivity and hyperresponsivity to suprathreshold sounds, particularly at the level of auditory cortex (Auerbach et al., 2014; Salvi et al., 2017; Herrmann and Butler, 2021a). In EEG/MEG recordings, hyperresponsivity is prominently represented as larger auditory cortex responses to sound onset for older compared to younger adults or for adults with hearing loss compared to adults without (Ross and Tremblay, 2009; Sörös et al., 2009; Lister et al., 2011; Alain et al., 2014; Bidelman et al., 2014; Parry et al., 2019; Irsik et al., 2021; Herrmann et al., 2022).

Consistent with hyperresponsivity to sound, neural synchronization with the low-frequency amplitude modulations of narrow-band sounds has been shown to be larger in older compared to younger people (Purcell et al., 2004; Goossens et al., 2016; Herrmann et al., 2019). Synchronization with the amplitude envelope of speech also appears to be greater in older compared to younger people (Presacco et al., 2016b, a; Decruy et al., 2020; Broderick et al., 2021) and in older adults with hearing loss compared to those without (Decruy et al., 2020; Fuglsang et al., 2020; but see Presacco et al., 2019). Synchronization seems particularly potentiate in older adults when the amplitude modulation envelope is characterized by rapid onsets and slow offsets compared to the reverse (Irsik et al., 2021). Age differences in neural synchronization to frequency-modulated, or amplitude- and frequency-modulated, sounds have been less investigated. In one study, neural synchronization in older people appeared to be increased when sounds were frequency modulated, consistent with hyperresponsivity (Boettcher et al., 2002). In another, no age difference in synchronization to a frequency-modulated sound was observed under passive listening conditions, whereas synchronization was reduced for older adults under active listening conditions (Henry et al., 2017). However, the sounds used in the Henry et al (2017) study contained short gaps that briefly interrupted the frequency modulation. Responses to gaps may differ between younger and older adults (Ross et al., 2010; Harris et al., 2012) and this may affect measured neural synchronization. Given this conflicting evidence, here we examine how the brains of younger (aged 21-33 years) and older (aged 53-73 years) people respond to sounds with low-frequency modulation in amplitude, frequency, and both combined.

Most studies that have investigated neural synchronization, and age differences therein, have focused on sinusoidal amplitude or frequency modulations (Boettcher et al., 2002; Goossens et al., 2016; Henry et al., 2017; Goossens et al., 2019; Herrmann et al., 2019) which are not very naturalistic. Natural speech contains low-frequency, non-sinusoidal fluctuations in amplitude and frequency; these include the amplitude changes associated with uttering syllables/words and frequency changes associated with the prosodic contour (Rosen, 1992; Varnet et al., 2017). Neural synchronization to the amplitude envelope of speech – reflecting non-sinusoidal amplitude fluctuations – has been shown to be greater for older compared to younger adults (Presacco et al., 2016b, a; Decruy et al., 2020; Broderick et al., 2021). Age differences in neural synchronization with non-sinusoidal frequency modulations have not yet been investigated.

Whether the sustained neural response differs between age groups has been investigated much less compared to age differences in neural synchronization. The existing work points to an age-related reduction in the sustained response. For example, short pure tones elicited a smaller sustained response in older compared to younger adults (Pfefferbaum et al., 1979). The sustained neural response elicited by repeated sets of tones in longer tone sequences has also been shown to be reduced for older relative to younger adults (Al Jaja et al., 2020; Herrmann et al., 2022). A similar age-related reduction in the sustained response has been reported in an EEG study for low-frequency amplitude modulations (Herrmann et al., 2019), although the effect of age group (younger vs older) was only marginally significant in this study. The sensitivity of MEG can be higher compared to EEG because skull and scalp tissue distort electric potentials more than magnetic fields (Hämäläinen et al., 1993; Hämäläinen and Hari, 2002). MEG may thus provide higher sensitivity to age-group differences. Moreover, whether the sustained response for frequency-modulated sounds or sounds with concurrent amplitude and frequency modulations – sinusoidal or non-sinusoidal – differs between younger and older adults has not been examined.

In the current study, MEG was recorded in two experimental sessions while younger and older adults listened to narrow-band noise sounds that were modulated in amplitude, frequency, or both concurrently. Amplitude and frequency were modulated either sinusoidally at 4 Hz or non-sinusoidally between 2 and 6 Hz. We examined the effects of modulation type (amplitude, frequency, both) and regularity (sinusoidal, non-sinusoidal) on neural synchronization and the sustained neural response, and whether these neural signatures differ between younger and older adults.

## Methods and Materials

### Participants

Twenty-six younger (mean: 26.7 years; range: 21–33 years; 13 males and 13 females) and twenty-five older adults (mean: 63.9 years; range: 53–73 years; 11 males and 14 females) were recruited for the current study. Participants provided written informed consent. None of the participants reported having a neurological disease, hearing impairment, or a hearing-aid prescription. Data were acquired in two sessions separated by at least one and maximal 43 days (median: 7 days). There was no age-group difference in the number of days between sessions (t49 = 0.99, p = 0.327). The study was conducted in accordance with the Declaration of Helsinki and the Canadian Tri-Council Policy Statement on Ethical Conduct for Research Involving Humans, and was approved by the Nonmedical Research Ethics Board of the University of Western Ontario.

Additional data (~12 min) for a different project (Herrmann et al., 2022) were recorded during each of the two ~1 hour sessions. Apart from audiometric hearing thresholds, none of the data reported here have been used or published previously.

## Audiometric hearing assessment and hearing thresholds

For each participant, pure-tone audiometric thresholds were estimated for pure tones at frequencies of 0.25, 0.5, 1, 2, 4, and 8 kHz (Figure 1). Mean thresholds averaged across 0.25, 0.5, 1, 2, and 4 kHz (i.e., pure-tone average) were higher for older compared to younger adults (t49 = 7.79, p = 4 × 10^−10^, d = 2.182; Figure 1, right; Herrmann et al., 2022). Although these thresholds are clinically ‘normal’ for age according to the ISO-7029 standard (https://www.iso.org/standard/42916.html), elevated thresholds are consistent with the presence of mild-to-moderate hearing loss in our sample of older adults, as would be expected (Moore, 2007; Plack, 2014; Presacco et al., 2016b; Herrmann et al., 2018).

**Figure 1:**
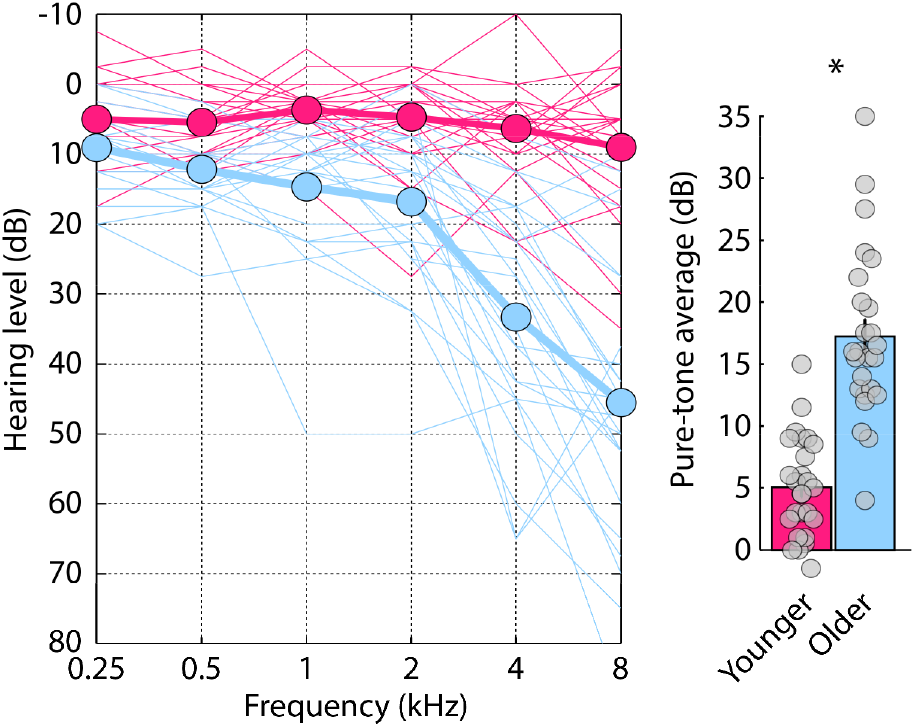
Audiograms and pure-tone average thresholds. Left: Pure-tone audiometric thresholds: The thin lines reflect data from each individual participant. Thick lines reflect the mean across participants. **Right:** Pure-tone average hearing threshold (mean across 0.25, 0.5, 1, 2, and 4 kHz). Gray dots reflect the pure-tone average threshold for individual participants. Error bars show the standard error of the mean. *p<0.05

In order to obtain a reference threshold in MATLAB software for sound presentation during the main experimental procedures, we also estimated the sensation level for a narrow-band noise that was generated similarly to the sounds used in the current study (see below), using a method-of-limits procedure (Herrmann et al., 2022). Participants listened to a 15-s narrow-band noise (~0.84–2.6 kHz) that changed continuously in intensity at a rate of 4 dB/s (either decreased [i.e., starting at suprathreshold levels] or increased [i.e., starting at subthreshold levels]). Participants pressed a button when they could no longer hear the tone (intensity decrease) or when they started to hear the tone (intensity increase); the sound stopped after the button press. The sound intensity at the time of the button press was noted for 6 decreasing sounds and 6 increasing sounds (decreasing and increasing sounds alternated), and these were averaged to determine the sensation level. As expected, given the audiometric pure-tone average thresholds (Figure 1), sensation levels were elevated for older compared to younger adults (t49 = 4.593, p = 3.07 × 10^−5^, d = 1.287, mean difference: 6.5 dB).

To approximately match audibility across age groups, all sound stimuli presented during the main experiment (described below) were presented at 55 dB above each individual’s sensation level. Since sensation levels and audiometric thresholds were on average larger for older compared to younger adults, sounds presented during the main experiment were on average more intense in sound-pressure level in older compared to younger adults. Previous research shows that brain responses to sound increase with increasing sound intensity (Picton et al., 1974; Picton et al., 1978; Pfefferbaum et al., 1979; Polich et al., 1988; Schadow et al., 2007; Herrmann et al., 2018). Because we predicted regularity-related sustained activity to be reduced in older compared to younger adults (Pfefferbaum et al., 1979; Al Jaja et al., 2020; Herrmann et al., 2022), the higher sound levels used with older adults works against this hypothesis. However, we are predicting that responses to sound onset and neural synchronization with temporal regularity in sounds to be larger in older compared to younger adults (Sörös et al., 2009; Alain et al., 2012; Goossens et al., 2016; Herrmann et al., 2019). A higher sound level could thus bias our data towards our hypothesized outcome. To mitigate this issue, data analysis for sound-onset responses and neural synchronization was conducted for a subset of 17 participants per age group for which sensation levels and sound-presentation levels did not statistically differ (mean difference: 1.09 dB; t32 = 1.331, p = 0.193, d = 0.456; for this, we selected the 17 younger adults with the highest sensation levels and the 17 older adults with the lowest sensation levels). Whereas the sensation levels and thus sound-presentation levels for the narrow-band noises did not differ between the younger and older sub-groups, pure-tone average thresholds were elevated for the older compared to the younger age sub-group (t32 = 5.825, p = 1.8 × 10^−6^, d = 1.998, mean difference: 7.77 dB; all <25 dB HL).

## Sound stimulation and procedure

All experimental procedures were carried out in an electromagnetically shielded, sound-attenuating room (Vacuumschmelze, Hanau). Sounds were presented binaurally via in-ear phones and the stimulation was controlled by a PC (Win XP, 64 Bit) running Psychtoolbox (v3.0.11) in Matlab (v7.8.0.347).

Acoustic stimuli were 4-s long narrow-band noises (see also Herrmann and Johnsrude, 2018; Herrmann et al., 2019). Narrow-band noises were created by adding 100 pure tones (components) with different carrier frequencies (Cf) that were randomly selected between 1200 and 2000 Hz, with random starting phase. Each of the 100 pure-tone components was amplitude and frequency modulated. The amplitude modulation depth was 100%. The range of the modulation in carrier frequency was defined as Cf ±Cf×0.3 (e.g., for a component with a mean carrier frequency of 1200 Hz, the carrier frequency modulation ranged between 840 Hz and 1560 Hz). The rate at which the amplitude and the carrier frequency was modulated was not fixed but changed randomly between 2 Hz and 6 Hz over time for each pure-tone component. The onset phase of the amplitude and frequency modulations as well as the randomization of amplitude- and frequency-modulation rate differed between components and between amplitude and frequency modulations. The 100 components were summed to obtain a narrow-band noise sound. We henceforth refer to this stimulus as the ‘random’ condition, because the signal resulting from the sum of the 100 pure-tone components does not contain any low-frequency (2–6 Hz) amplitude or frequency modulations (Figure 2 top).

**Figure 2:**
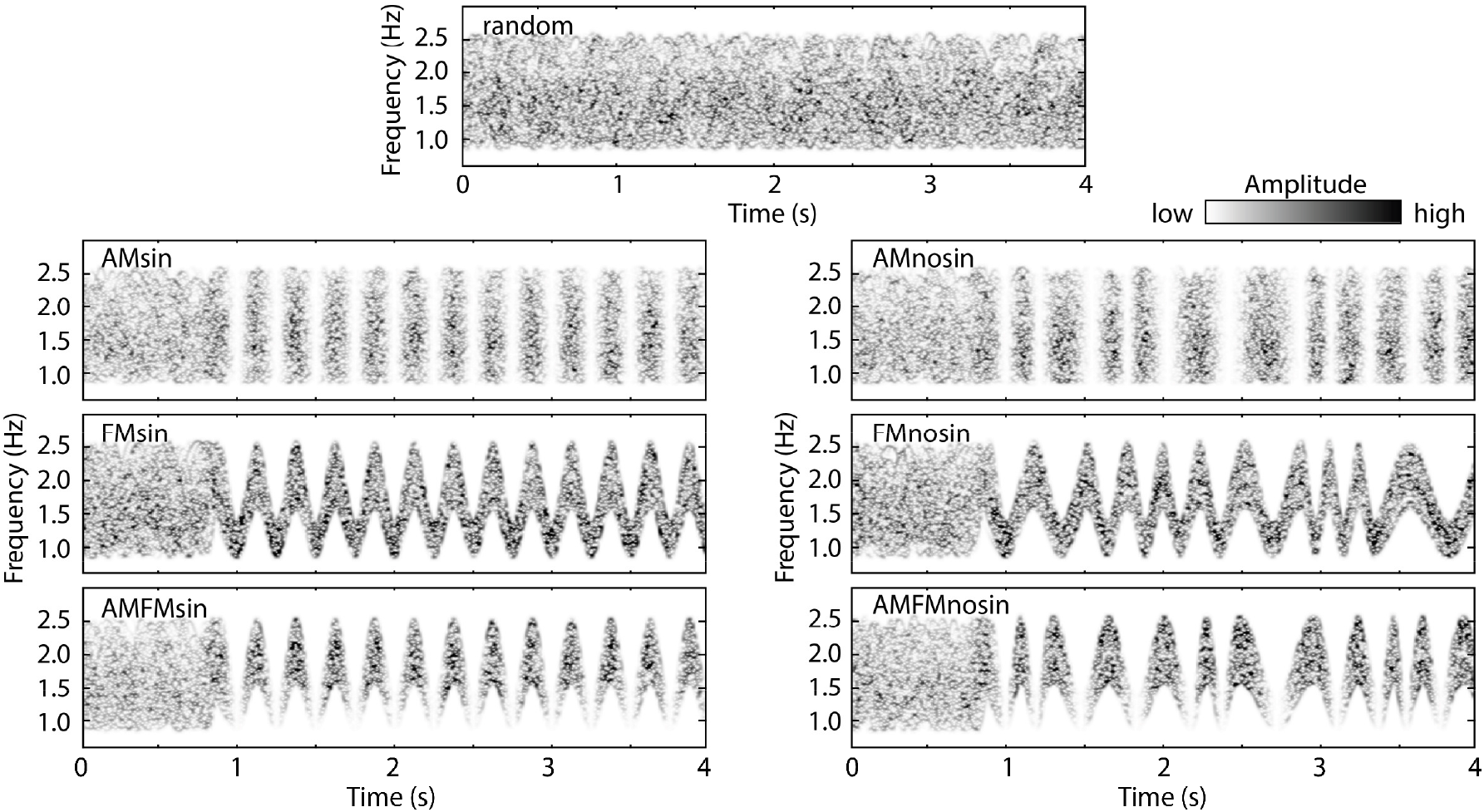
Stimulus conditions. Spectrograms for samples of narrow-band noise sounds for each of the seven stimulus conditions. The ‘random’ condition did not contain low-frequency amplitude or frequency modulations and had a relatively consistent spectrum over time. The other six conditions smoothly transitioned to sinusoidally (left) or non-sinusoidally (right) amplitude, frequency, or amplitude and frequency-modulated noises at around 1 s after sound onset. Abbreviations: sinusoidal amplitude modulation (AMsin), sinusoidal frequency modulation (FMsin), sinusoidal amplitude and frequency modulation (AMFMsin), non-sinusoidal amplitude modulation (AMnosin), non-sinusoidal frequency modulation (FMnosin), non-sinusoidal amplitude and frequency modulation (AMFMnosin).

Six additional conditions were created in which either the amplitude modulation (AM), frequency modulation (FM), or both the amplitude and frequency modulations (AMFM) of each of the 100 pure-tone components became synchronized (phase-aligned) approximately 1 second after sound onset. There was no transient change at the transition from non-aligned to AM/FM phase-aligned pure-tone components (see also Herrmann and Johnsrude, 2018; Herrmann et al., 2019; Figure 2). The modulation rate of the AM, FM, or AMFM after phase-alignment was either constant and sinusoidal at 4 Hz (Figure 2 left) or changed randomly and non-sinusoidally between 2 and 6 Hz for the rest of the sound (from ~1 s to 4 s; Figure 2 right). Averaging the 100 pure-tone components resulted in a narrow-band noise sound that was non-modulated (random) for about 1 second, after which it transitioned to a sinusoidal or non-sinusoidal AM, FM, or AMFM. Henceforth, we refer to the six conditions as ‘sinusoidal amplitude modulation’ (AMsin), ‘sinusoidal frequency modulation’ (FMsin), ‘sinusoidal amplitude and frequency modulation’ (AMFMsin), ‘non-sinusoidal amplitude modulation’ (AMnosin), ‘non-sinusoidal frequency modulation’ (FMnosin), ‘non-sinusoidal amplitude and frequency modulation’ (AMFMnosin).

In each of the two recording sessions, participants were presented with five ~10-min blocks of stimulation. Participants listened passively to 70 trials of each condition per session (5 blocks × 14 trials per condition = 70 trials), and thus to 140 trial per condition across the study. Trials of the seven conditions were presented pseudo-randomly throughout a block, such that each condition could occur maximally three times in direct succession. Trials were separated by a 2-s inter-stimulus interval. During sound presentation, participants watched a muted, subtitled, movie of their choice that was projected into the electromagnetically shielded room via a mirror system.

### Recordings and initial preprocessing of magnetoencephalographic data

Methods of data recording and preprocessing were similar to our previous work (Herrmann et al., 2022). For each participant, magnetoencephalography was recorded in an electromagnetically shielded room (AK3b, Vacuumschmelze, Hanau, Germany) using a 306-channel Neuromag Vectorview MEG (MEGIN Oy, Helsinki, Finland) at the Max Planck Institute for Human Cognitive and Brain Sciences in Leipzig, Germany. Data were recorded with a sampling rate of 1000 Hz and an online low-pass filter of 330 Hz (no online high-pass filter). The signal space separation (SSS) method (maxfilter© version 2.2.15; default parameter setting Lin = 8; Lout = 3) was used to suppress external interference, interpolate bad channels (identified through visual inspection), and transform each person’s individual data to the sensor space of the first block of the first session to ensure the data are in a common space (Taulu et al., 2004; Taulu et al., 2005).

### Combination of magnetometer and gradiometer channels

Magnetic fields were recorded by one magnetometer channel and two gradiometer channels in each of 102 spatial locations distributed around a participant’s head. Magnetometers measure magnetic fields in Tesla (T), whereas gradiometers (a coupled pair of magnetometers) measure differences in the same magnetic fields over a distance of 0.0168 m in Tesla per meter (T/m). Combining the two MEG channel types for data analysis requires accounting for their different units. All gradiometer channels were transformed to magnetometer channels, because this transformation only requires a linear interpolation that results in the same unit for all channels (Herrmann et al., 2018, 2022). To this end, a scaling matrix **S** was multiplied with the data matrix **D** to obtain scaled data according to the following equation:

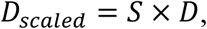

where **D** is a 3 × n matrix. The three rows of **D** refer to the two gradiometers and one magnetometer of one MEG sensor triplet (i.e., in one spatial location) and n to the number of data samples over time. The scaling matrix **S** is a 5 × 3 matrix with the following elements:

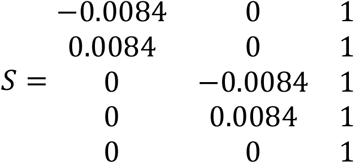

The value 0.0084 reflects half of the distance between the two gradiometer loops measured in meters, and the transformation constitutes a linear approximation of the magnetic field at each of the triplets. The transformation replaces the sensor triplet by a sensor quintet of magnetometers. The columns of **S** refer to the triplet of two gradiometers and one magnetometer and the rows of **S** refer to the resulting five magnetometers. This procedure resulted in signals from 510 magnetometer channels centered on and around 102 locations around a participant’s head (Herrmann et al., 2018, 2022).

### Preprocessing of magnetoencephalographic data

A high-pass (0.7 Hz; 2391 points, Hann window) and a low-pass filter (20.3 Hz, 119 points, Kaiser window) were applied to the data. Data were downsampled from the 1000-Hz to a 250-Hz sampling frequency and epochs ranging from −1 s pre-stimulus onset to 5 s post-stimulus onset were extracted. Independent components analysis was calculated for epoched data using the Fieldtrip MATLAB toolbox (v20130727, Oostenveld et al., 2011) with the runica method (Makeig et al., 1996) and the logistic infomax algorithm (Bell and Sejnowski, 1995) to identify and remove activity related to blinks, horizontal eye movements, muscle activity, and noisy channels. Finally, epochs that contained a signal change larger than 8 picotesla (pT) in any channel were excluded. These data were used to investigate condition and age differences in evoked responses to sound onset and neural synchronization.

The same analysis pipeline was separately run with slightly different parameters to investigate the sustained response. The sustained neural response is a very low-frequency signal reflecting an offset shift (Barascud et al., 2016; Southwell et al., 2017; Herrmann and Johnsrude, 2018). Application of a high-pass filter would remove this signal and was thus omitted for the analysis of the sustained response. Moreover, sounds contained temporal modulations (AM, FM, AMFM) at 4 Hz (sinusoidal) and 2–6 Hz (non-sinusoidal) that drive neural activity in the corresponding neural frequency bands. In order to avoid that the measured sustained response is influenced by neural-activity changes in the 2–6 Hz frequency band, data were filtered with a 1.2 Hz low-pass filter (3001 points, Kaiser window). The analysis pipeline was otherwise similar. The 0-1.2 Hz band data were downsampled to 250 Hz and divided into 6-s epochs. Activity related to blinks, horizontal eye movements, muscle activity, and noisy channels was removed using the identified components from the 0.7-20.3 Hz band data. Epochs in which a signal change larger than 8 pT occurred in any channel were excluded.

### Analysis of responses to sound onset

Responses to sound onset were analyzed using the 0.7-20.3 Hz band data. To investigate whether older adults exhibit hyperresponsivity to sound, and thus possibly a loss of inhibition, we examine the M50 and M100 responses to sound onset (cf. Sörös et al., 2009; Alain et al., 2012; Herrmann et al., 2018). Sounds from different conditions were similar for the first second after sound onset, and data for this onset period were thus averaged across all epochs (ignoring conditions). Absolute values were calculated for signals of each channel to account for opposite polarities of the magnetic fields in directions perpendicular to the tangential orientation of the underlying neural source. Separately for each channel, the mean signal from the 0.2 s period immediately prior to sound onset was subtracted from the signal at each time point (baseline correction). In order to focus the analysis on the most sensitive channels, signals were averaged across sensors located over temporal cortex (Figure 3). This resulted in one response time course per participant and hemisphere (left, right).

**Figure 3:**
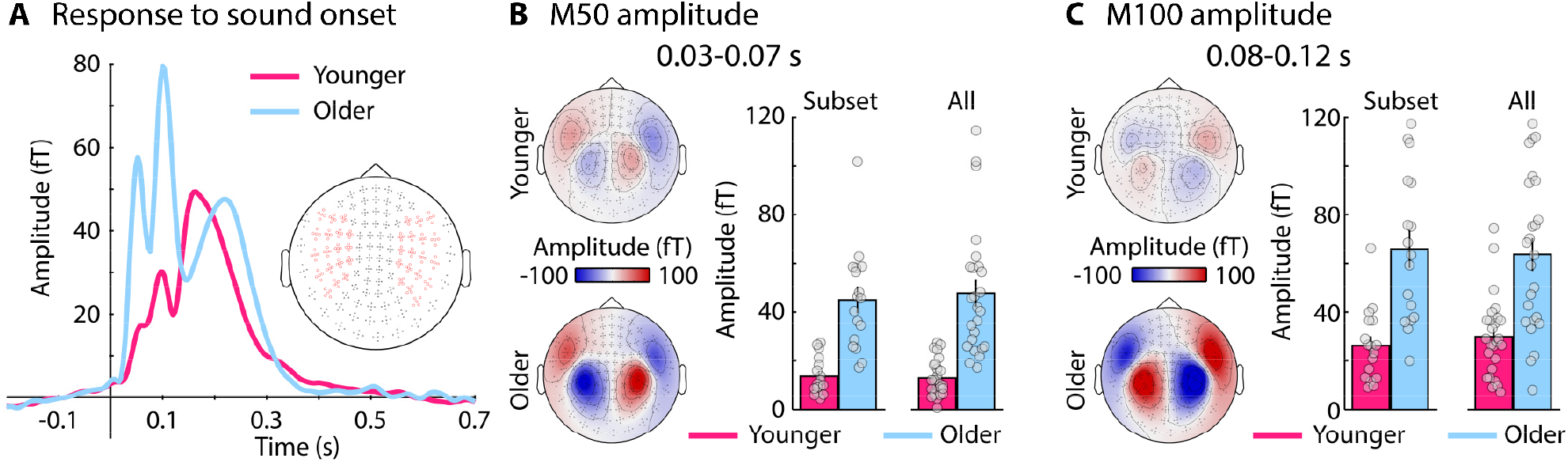
Results for sound-onset responses. **A:** Time courses in response to sound onset (averaged across the left and right temporal-cortex channels). Temporal-cortex channels are marked in red in the channel display. **B:** M50 topographies and mean responses (0.03–0.07 s). **C:** M100 topographies and mean responses (0.8–0.12 s). Gray dots in panels B and C reflect data from individual participants. Data in panel A and topographies in panels B and C reflect the averaged data of the seventeen participants in each age group for which sensation levels did not differ statistically. Error bars reflect the standard error of the mean.

For the statistical analysis, responses were separately averaged in the 0.03–0.07 s and the 0.08–0.12 s time windows for the M50 and M100, respectively. Separate repeated measures analyses of variance (rmANOVA) were calculated for the M50 and M100 responses, comprising the within-participants factor Hemisphere (left, right) and the between-participants factor Age Group (younger, older). We focused analyses on the 17 participants from each age group for which sensation level – and thus the level at which sounds were delivered – did not differ between age groups, but data for all participants are provided as well.

### Analysis of neural synchronization: Cross-coherence

Neural synchronization was analyzed using the cross-coherence measure (Nolte et al., 2004; Keitel et al., 2016). Cross-coherence is related to the more common measures of neural synchronization, namely intertrial phase coherence or evoked amplitude/power (Lachaux et al., 1999; Picton et al., 2003; Nozaradan et al., 2011; Anderson et al., 2012; Herrmann et al., 2013; Herrmann and Johnsrude, 2018). Inter-trial phase coherence and evoked amplitude/power require that neural activity is temporally consistent across trials. In the current study, the temporal modulations of sounds with a non-sinusoidal stimulus rhythm differed from trial to trial. Inter-trial phase coherence and evoked amplitude/power are thus not suitable for the analysis of neural activity in the current study. Cross-coherence provides an alternative that is similarly interpreted as inter-trial phase coherence (Nolte et al., 2004; Keitel et al., 2016).

Calculation of cross coherence focused on the 1–4 s time window following sound onset (the first second did not contain any low-frequency AM, FM, or AMFM) and involved the following steps. A fast Fourier transform (FFT; Hann window; zero-padding) was calculated for each single-trial stimulus time course and for each single-trial neural time course (separately for each MEG channel). This resulted in a stimulus and a neural spectrum of complex numbers for each trial and channel. For each MEG channel and neural-frequency bin in the FFT spectrum, coherence was then calculated between the stimulus spectrum and the neural spectrum according to the following equation (Nolte et al., 2004):

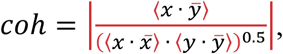

where x and y reflect the complex numbers and 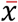 and 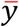 the complex conjugates of the stimulus and the neural spectrum, respectively. The dot refers to the multiplication operation. The bracket notation ⟨ ⟩ refers to the expected value, that is, the mean across trials. The notation | | reflects the magnitude of the complex number.

To analyze cross-coherence for the 4-Hz sinusoidal stimulation, cross-coherence was averaged in the 3.95 to 4.05 Hz frequency window. To analyze cross-coherence for the 2–6 Hz non-sinusoidal stimulation, coherence was averaged in the 2–6 Hz frequency window. Cross-coherence was averaged across temporal-cortex channels.

For the statistical analyses, cross-coherence for the random condition was subtracted from the cross-coherence of each of the other six conditions (AMsin, FMsin, AMFMsin, AMnosin, FMnosin, AMFMnosin). Statistical analyses were conducted separately for sinusoidal and non-sinusoidal stimulus conditions, because the different stimulation frequencies resulted in qualitatively different coherence spectra (see Figure 4). Hence, two separate repeated-measures ANOVAs were calculated (sinusoidal; non-sinusoidal) using cross-coherence as the dependent measure. The within-participants factors were Modulation Type (AM, FM, AMFM) and Hemisphere (left, right), and the between-participants factor was Age Group (younger, older). The Greenhouse and Geisser correction was used when Mauchly’s test indicated that sphericity was violated (Greenhouse and Geisser, 1959). The original degrees of freedom are reported, along with the epsilon coefficient, and the corrected p-value. Analyses focused on the 17 participants from each age group for which sensation level did not differ between age groups, but data for all participants are provided as well.

**Figure 4:**
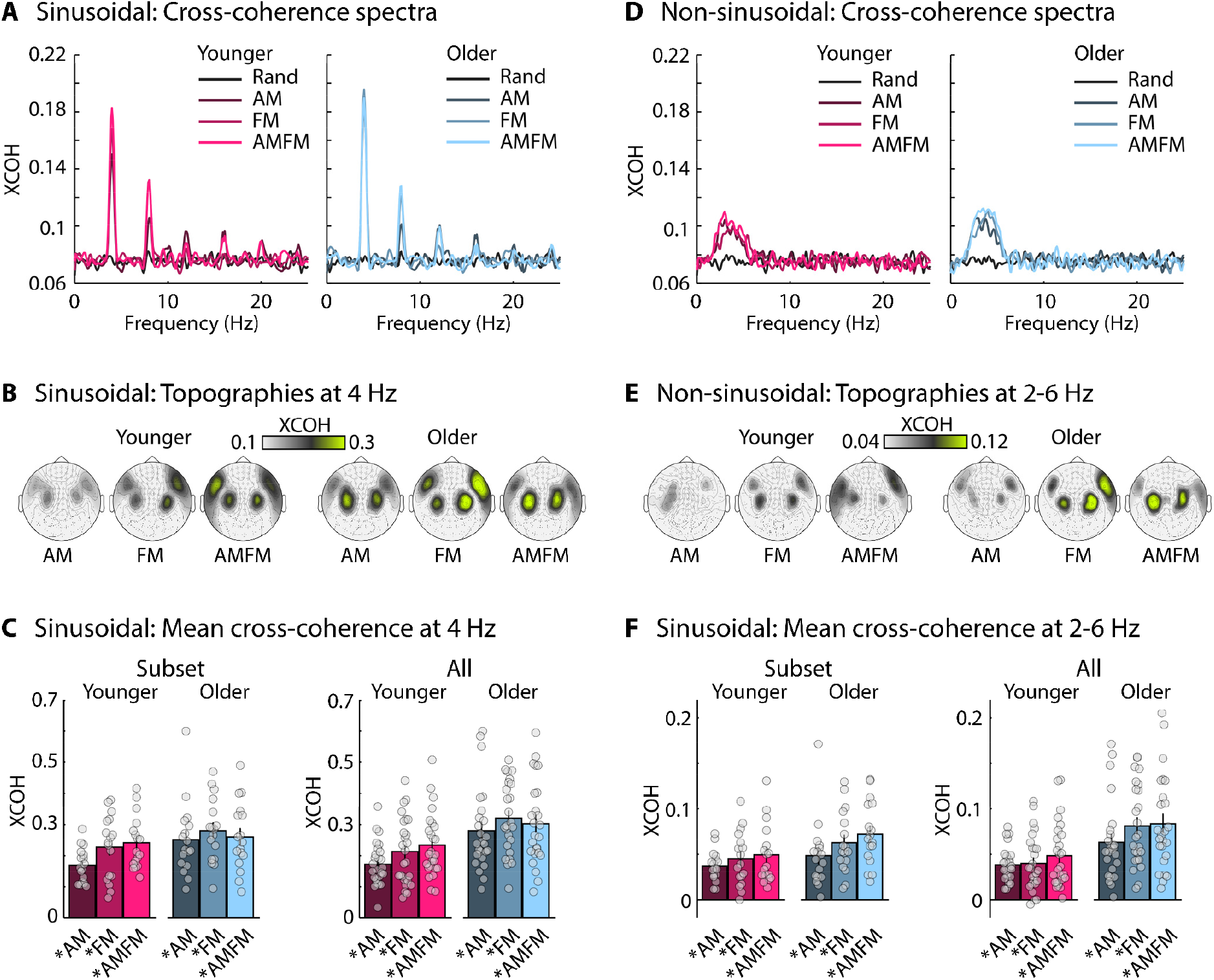
Results for the cross-coherence (XCOH) analysis. **A:** Cross-coherence spectra for sinusoidal sound stimulation (averaged across the left and right temporal-cortex channels). **B:** Topographical distributions for cross-coherence at 4 Hz for sinusoidal stimulation. **C:** Mean cross-coherence at 4 Hz for sinusoidal stimulation. The asterisk indicates a significant difference between the respective condition relative to the random condition (p ≤ 0.05; FDR-corrected). **D-F:** Same as for panels A-C for non-sinusoidal sound stimulation and 2–6 Hz cross-coherence. Gray dots in panels C and F reflect data from individual participants. Data in panels A, B, D, and E reflect the averaged data of the seventeen participants in each age group for which sensation levels did not differ statistically. Error bars reflect the standard error of the mean.

### Analysis of neural synchronization: Relationship between stimulus phase and neural response

Cross-coherence provides a spectrum that can be interpreted similar to inter-trial phase coherence (Lachaux et al., 1999; Nolte et al., 2004; Keitel et al., 2016). However, non-sinusoidal responses in the time domain lead to distributed energy across the fundamental frequency and its harmonics in the cross-coherence spectrum (due to the sinusoidal assumption of the Fourier transform). Moreover, cross-coherence spectra cannot be compared between sinusoidal and non-sinusoidal modulations due to their different characteristics. To account for these disadvantages associated with coherence spectra, we analyzed neural response amplitude as a function of stimulus phase (Irsik et al., 2021). To this end, the stimulus AM, FM, or AMFM of each trial was extracted from the 1–4 s time window after stimulus onset and downsampled to match the sampling of the MEG signal. Because the random condition did not contain a coherent low-frequency temporal modulation, we randomly selected, for each random-condition trial, a stimulus AM, FM, or AMFM from another trial within the same block. The analytic Hilbert transform was calculated for each trial-specific stimulus modulation (AM, FM, or AMFM), and the resulting complex numbers were used to calculate the phase angle for each time sample within 1–4 s. As a result, data for each trial comprised a stimulus phase series and the corresponding neural amplitudes (for each MEG channel). The neural amplitude data were binned according to the stimulus phase values (number of bins: 100; window width: 0.314 radians; with overlap), such that neural amplitude was represented as a function of stimulus phase ranging from −π to π (binning accounted for circularity). This resulted in a phase-amplitude series for each MEG channel, reflecting the neural response in stimulation cycles.

In the phase-amplitude series, the amplitude range between positive and negative deflections captures the magnitude of the synchronized response. In order to calculate whether neural responses differed between conditions and age groups, the amplitude range was calculated separately for each stimulus condition (random, AMsin, FMsin, AMFMsin, AMnosin, FMnosin, AMFMnosin), age group (younger, older), and MEG channel as the difference between the maximum and the minimum amplitude in the phaseamplitude series. The amplitude range was averaged across temporal-cortex channels.

For statistical analyses, the amplitude range of the random condition was separately subtracted from the amplitude range of each of the other six conditions (AMsin, FMsin, AMFMsin, AMnosin, FMnosin, AMFMnosin). A rmANOVA was calculated using amplitude range as the dependent measure. Within-participants factors were Modulation Type (AM, FM, AMFM), Regularity (sinusoidal, non-sinusoidal), and Hemisphere (left, right), and the between-participants factor was Age Group (younger, older). The Greenhouse and Geisser correction was used when Mauchly’s test indicated that sphericity was violated (Greenhouse and Geisser, 1959). The original degrees of freedom are reported, along with the epsilon coefficient, and the corrected p-value. Analyses focused on the 17 participants from each age group for which sensation level did not differ between age groups, but bar graphs and individual data points are also provided.

### Analysis of the sustained response

The sustained neural responses were analyzed using the 0-1.2 Hz band data. Single-trial time courses ranging from −1 to 5 s time-locked to sound onset were averaged, separately for each condition (random, AMsin, FMsin, AMFMsin, AMnosin, FMnosin, AMFMnosin). The absolute values of the averaged time courses were calculated for each channel. The mean signal in the 1-s time window prior to sound onset was averaged and subtracted from the signal at each time point, separately for each channel (baseline correction). Signals were averaged across the temporal-cortex channels, resulting in one response time course per condition, hemisphere, and participant.

For statistical analyses, responses for the random condition were subtracted from the responses of each of the other six conditions. Statistical comparisons focused on the 2–3.5 s time window. The start of the time window at 2 s is similar to our previous work were sounds changed to a regular pattern approximately 1 s after sound onset (Herrmann et al., 2022) and allows for the auditory system to be fully responsive to the temporal modulations (see also Barascud et al., 2016; Teki et al., 2016; Herrmann and Johnsrude, 2018). The very low-frequency low-pass filter can potentially smear neural responses in time. To reduce the possibility that the response to the sound offset at 4 s affects our measurement of the sustained response, we chose to limit the analysis time window to 3.5 s instead 4 s.

An rmANOVA was calculated using the sustained response as the dependent measure. Modulation Type (AM, FM, AMFM), Regularity (sinusoidal, non-sinusoidal), and Hemisphere (left, right) were within-participants factors, and Age Group (younger, older) was a between-participants factor. The Greenhouse-Geisser correction was used when Mauchly’s test indicated that sphericity was violated (Greenhouse and Geisser, 1959). The original degrees of freedom, epsilon coefficient, and corrected p-value are reported. All participants were included in the analysis of the sustained response, since the more intense stimuli presented to older people would lead to larger responses, if anything, and we hypothesized that sustained activity would be reduced in older compared to younger adults (Pfefferbaum et al., 1979; Al Jaja et al., 2020; Herrmann et al., 2022).

## Results

### Increased response to sound onset in older adults

Time courses of neural responses to sound onset are displayed in Figure 3A. Responses were larger for older compared to younger adults for both the M50 time window (0.03–0.07 s: F1,32 = 34.109, p = 1.7 · 10^−6^, ω^2^ = 0.334; Figure 3B) and the M100 time window (0.08–0.12 s: F1,32 = 23.176, p = 3.4 · 10^−5^, ω^2^ = 0.251; Figure 3C). There was no difference between hemispheres nor an Age Group × Hemisphere interaction, for neither time window (p > 0.05). This is consistent with previous work showing hyperresponsivity to sound in the auditory cortex of older compared to younger adults (Anderer et al., 1996; Sörös et al., 2009; Alain et al., 2012; Herrmann et al., 2022). Topographical distributions indicate neural generators in auditory cortex (Herrmann et al., 2022).

### Increased neural synchronization in older adults

Neural synchronization was first analyzed using cross-coherence. Repeated-measures ANOVAs were conducted separately for stimuli with sinusoidal and non-sinusoidal modulations, because different frequency bands were used to capture synchronized activity: 4 Hz for sinusoidal stimulation and 2–6 Hz for non-sinusoidal stimulation, corresponding to the respective stimulation frequencies.

Cross-coherence for 4-Hz sinusoidally modulated sounds did not differ between age groups (effect of Age Group: F1,32 = 2.482, p = 0.125, ω^2^ = 0.022). Across both age groups, cross-coherence was larger for FM (F1,33 = 4.669, p = 0.038, ω^2^ = 0.023) and AMFM sounds (F1,33 = 6.414, p = 0.016, ω^2^ = 0.035) compared to AM sounds (effect of Modulation Type: F2,64 = 3.906, p = 0.025, ω^2^ = 0.026), but coherence for FM and AMFM did not differ (F1,33 = 0.138, p = 0.713, ω^2^ < 0.001; Figure 4A-C). Other main effects and interactions were not significant. Results for sounds with a 2–6 Hz non-sinusoidal modulation showed larger, but only marginally significant cross-coherence for older compared to younger adults (effect of Age Group: F1,32 = 3.260, p = 0.080, ω^2^ = 0.033). Cross-coherence was larger for AMFM compared to AM sounds (F1,33 = 7.325, p = 0.011, ω^2^ = 0.045; effect of Modulation Type: F2,64 = 4.000, p = 0.023, ω^2^ = 0.03), but did not differ between AM and FM (F1,33 = 1.659, p = 0.207, ω^2^ = 0.004) nor between FM and AMFM (F1,33 = 2.721, p = 0.109, ω^2^ = 0.012; Figure 4D-F). Other main effects and interactions were not significant.

Neural synchronization was also quantified by calculating the relation between stimulus phase and neural response amplitude. The rmANOVA for the amplitude range in the phase-amplitude response curves showed larger response ranges for FMs than AMs (F1,33 = 5.159, p = 0.030, ω^2^ = 0.023), for AMFMs than AMs (F1,33 = 32.006, p = 2.65 · 10^−6^, ω^2^ = 0.12), and for AMFMs than FMs (F1,33 = 9.540, p = 0.004, ω^2^ = 0.043; effect of Modulation Type: F2,64 = 15.265, p = 3.8 · 10^−6^, ω^2^ = 0.097). Sinusoidal stimulation also led to larger amplitude ranges compared to non-sinusoidal stimulation (effect of Regularity: F1,32 = 103.625, p = 1.46 · 10^−11^, ω^2^ = 0.202). The Modulation Type × Regularity interaction was significant (F2,64 = 3.586, p = 0.033, ω^2^ = 0.007). We confirmed for each of the different modulation types (AM, FM, AMFM) that the sinusoidally modulated sounds elicited larger amplitude ranges than non-sinusoidally modulated sounds (for all: F1,33 > 30, p < 0.001, ω^2^ > 0.09). The amplitude range was larger in the left than the right hemisphere (effect of Hemisphere: F1,32 = 14.707, p = 5.6 · 10^−4^, ω^2^ = 0.078).

Critically, we observed larger amplitude ranges for older compared to younger adults (effect of Age Group: F1,32 = 5.305, p = 0.028, ω^2^ = 0.061), but no interactions involving Age Group (p > 0.05). However, given that the Regularity × Age Group interaction was marginally significant (F1,32 = 3.658, p = 0.065, ω^2^ = 0.007), we confirmed that larger amplitude ranges for older compared to younger adults were observed for sinusoidal (F1,32 = 5.436, p = 0.026, ω^2^ = 0.063) and non-sinusoidal sound stimulation (F1,32 = 4.563, p = 0.04, ω^2^ = 0.051; Figure 5).

**Figure 5:**
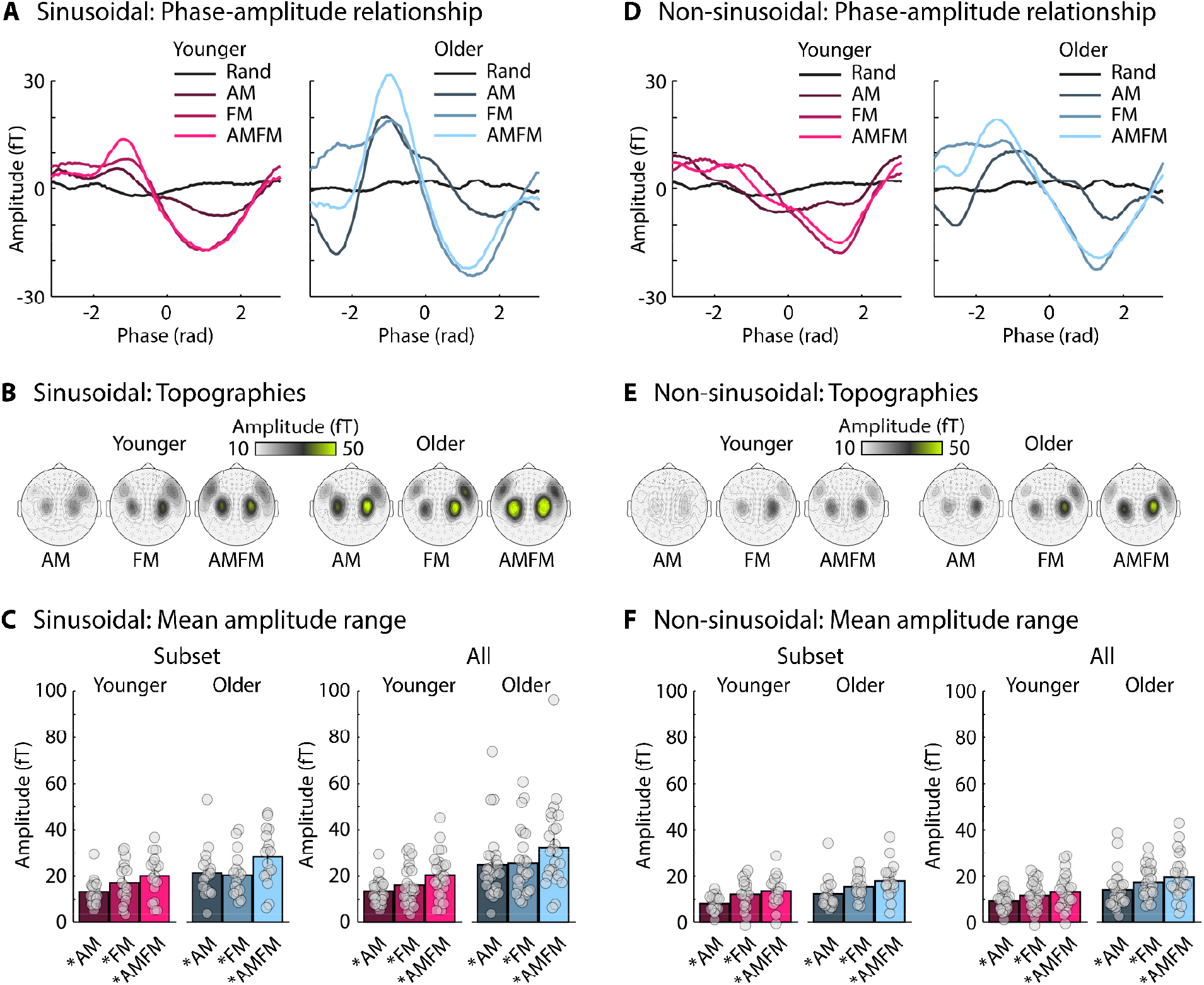
Results for the phase-amplitude analysis. **A:** Neural amplitude as a function of stimulus phase for sinusoidal sound stimulation (averaged across the left and right temporal-cortex channels). **B:** Topographical distributions for amplitude ranges for sinusoidal stimulation. **C:** Mean amplitude range for sinusoidal stimulation. The asterisk indicates a significant difference between the respective condition relative to the random condition (p ≤ 0.05; FDR-corrected). **D-F:** Same as for panels A-C for non-sinusoidal sound stimulation. Gray dots in panels C and F reflect data from individual participants. Data in panels A, B, D, and E reflect the averaged data of the seventeen participants in each age group for which sensation levels did not differ statistically. Error bars reflect the standard error of the mean.

Topographical distributions of cross-coherence (Figure 4B, 4E) and responses in phase-amplitude curves (Figure 5B, 5E) suggest that auditory cortex underlies neural synchronization activity.

### Reduced sustained neural response in older adults

Response time courses for sustained neural responses are displayed in Figure 6A. The mean sustained response in the 2–3.5-s time window was analyzed using an rmANOVA. The sustained response was larger for FMs (F1,50 = 35.797, p = 2.33 · 10^−7^, ω^2^ = 0.132) and AMFMs (F1,50 = 49.253, p = 5.623 · 10^−9^, ω^2^ = 0.146) compared to AMs (effect of Modulation Type: F2,98 = 35.711, p = 1.06 · 10^−10^, ε = 0.834, ω^2^ = 0.131), but did not differ between FM and AMFM (F1,50 = 0.102, p = 0.751, ω^2^ < 0.001). The sustained response was larger for non-sinusoidal compared to sinusoidal sound stimulation (effect of Regularity: F1,49 = 5.146, p = 0.028, ω^2^ = 0.024), and larger in the right compared to left hemisphere (effect of Hemisphere: F1,49 = 7.156, p = 0.01, ω^2^ = 0.008).

**Figure 6:**
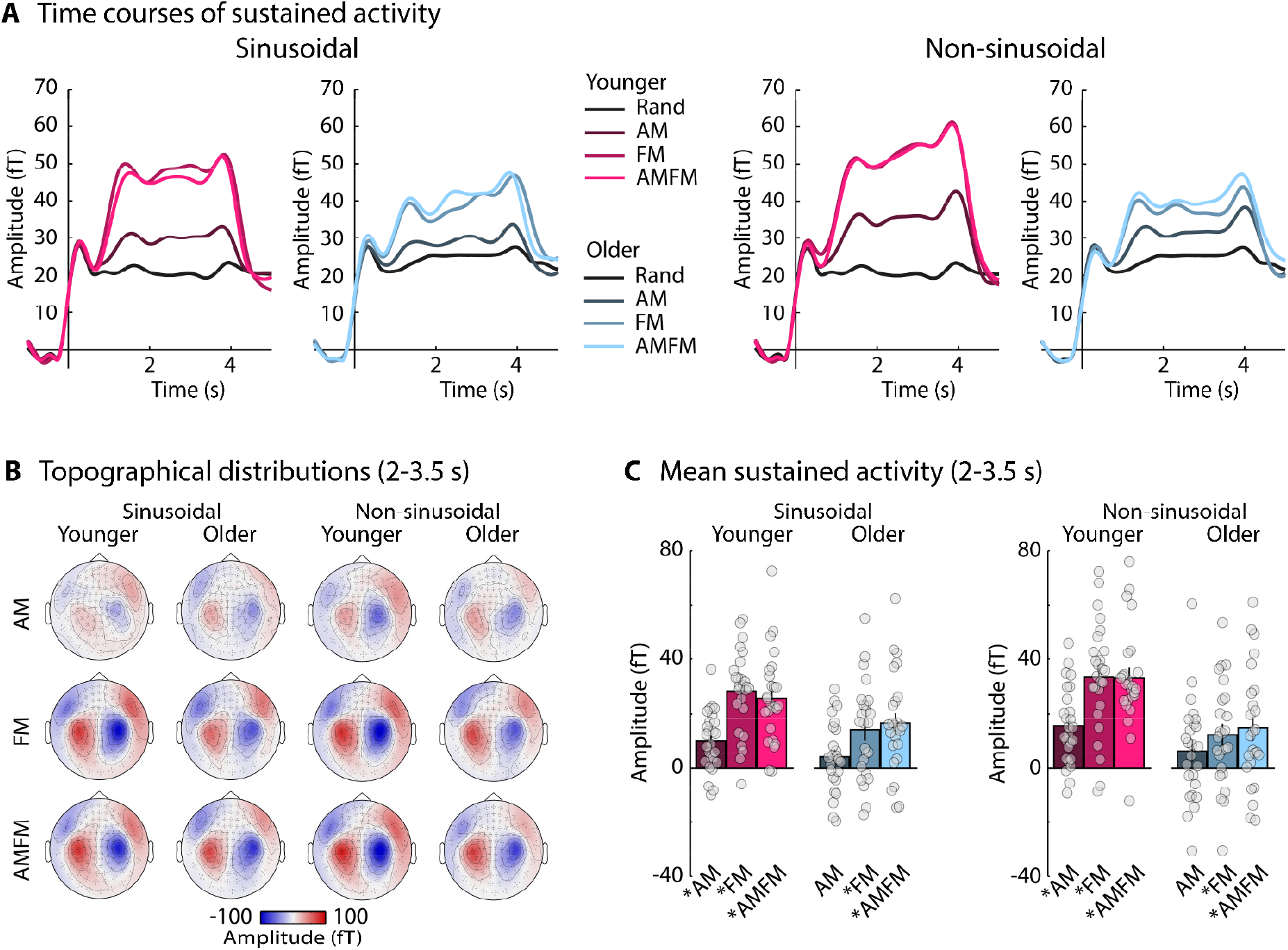
Results for the sustained response. **A:** Time courses for sinusoidal and non-sinusoidal sound stimulation (averaged across the left and right temporal-cortex channels). **B:** Topographical distributions for mean sustained activity in the 2–3.5-s time window. **C:** Mean sustained activity in the 2–3.5-s time window. The asterisk indicates a significant difference between the respective condition and the random condition (p ≤ 0.05; FDR-corrected). Error bars reflect the standard error of the mean. Gray dots in panels C reflect data from individual participants.

Critically, the sustained response was larger for younger compared to older adults (effect of Age Group: F1,49 = 10.885, p = 0.002, ω^2^ = 0.09). The Modulation Type × Age Group interaction was significant (F2,98 = 4.127, p = 0.026, ε = 0.946, ω^2^ = 0.013), but the age-group difference was present for AM (F1,49 = 4.388, p = 0.041, ω^2^ = 0.033), FM (F1,49 = 13.644, p = 5.56 · 10^−4^, ω^2^ = 0.112), and AMFM stimulation (F1,49 = 8.03, p = 0.007, ω^2^ = 0.066; Figure 6). The difference between age groups was larger for non-sinusoidal compared to sinusoidal stimulation (Regularity × Age Group interaction: F1,49 = 9.734, p = 0.003, ω^2^ = 0.011), but the effect of Age Group was significant for both (sinusoidal: F1,49 = 7.176, p = 0.01, ω^2^ = 0.058; non-sinusoidal: F1,49 = 13.249, p = 6.56 × 10^−4^, ω^2^ = 0.109). Moreover, the increase of the sustained response for non-sinusoidal compared to sinusoidal modulations was only significant for younger (F1,25 = 15.548, p = 5.73 · 10^−4^, ω^2^ = 0.048), but not for older adults (F1,24 = 0.108, p = 0.745, ω^2^ < 0.001). Other main effects and interactions were not significant.

## Discussion

The current study investigated whether neural synchronization and sustained responses to low-frequency modulations in a sound’s amplitude, frequency, or both differ between younger and older adults. The MEG data demonstrate that neural synchronization was reduced for non-sinusoidal compared to sinusoidal modulation, but both were stronger, across all modulation types, in older compared to younger adults. The sustained neural response elicited by the frequency and combined amplitude-frequency modulations was reduced for older compared to younger adults. The sustained response was also reduced for sinusoidal compared to non-sinusoidal sound stimulation in younger, but not in older adults. The results indicate that the auditory systems of older and younger adults differ in their response to amplitude, frequency, and combined amplitude-frequency modulations in sounds.

### Neural activity is sensitive to low-frequency modulations in amplitude, frequency, and both combined

Synchronization with low-frequency amplitude and frequency modulations has been reported previously (Maiste and Picton, 1989; Boettcher et al., 2002; Picton et al., 2003; Goossens et al., 2016; Henry et al., 2017; Irsik et al., 2021), although not all studies focus on this speech-relevant frequency region of ~4 Hz (Edwards and Chang, 2013). Here, neural synchronization was investigated using cross-coherence and a phase-amplitude analysis. Cross-coherence (Nolte et al., 2004) is an alternative metric to inter-trial phase coherence and allows the investigation of neural synchronization to non-sinusoidal modulations (Keitel et al., 2016). Frequency modulations and amplitude-frequency modulations led to larger synchronization than did amplitude modulations (Figure 4). The phase-amplitude analysis (Irsik et al., 2021) appeared to be somewhat more sensitive, showing a larger response for combined amplitude-frequency modulations compared to amplitude or frequency modulation alone (Figure 5). In the current study amplitude and frequency modulation were not perceptually matched in, for example, perceived fluctuation strength (Fastl, 1982; Fastl, 1983; Moore et al., 1996; Fastl and Zwicker, 2007; Vecchi et al., 2016). Previous research suggests that perceived fluctuation strength for AMs is lower compared to FMs with a modulation depth similar to the one used here (Fastl and Zwicker, 2007; Vecchi et al., 2016). Hence, the frequency modulation may have been perceived as fluctuating more strongly than the amplitude modulation, potentially explaining the larger synchronization.

The sustained response was also sensitive to amplitude and frequency modulations. This is consistent with previous work demonstrating a sustained response to low-frequency amplitude and frequency modulations in sounds (Herrmann and Johnsrude, 2018; Herrmann et al., 2019), and more generally with an increasing body of work showing that temporally regular patterns elicit a sustained neural response (Barascud et al., 2016; Sohoglu and Chait, 2016; Teki et al., 2016; Southwell et al., 2017; Southwell and Chait, 2018; Dheerendra et al., 2021; Herrmann et al., 2021). Similar to neural synchronization, the sustained response was larger for frequency modulations and amplitude-frequency modulations compared to amplitude modulations (Figure 6). This suggests that frequency modulations compared to amplitude modulations are especially potent drivers of the sustained response and, as mentioned above, this may reflect perceived fluctuation strength.

### Sinusoidal vs non-sinusoidal temporal modulations differently affect neural activity

Quasi-rhythmic stimulation has been shown to drive synchronized activity (Stefanics et al., 2010; Keitel et al., 2016), including the tracking of non-sinusoidal amplitude fluctuations in speech (Giraud and Poeppel, 2012; Obleser et al., 2012; Peelle and Davis, 2013; Peelle et al., 2013; Brodbeck and Simon, 2020). Consistent with this, we observed that neural synchronization was significant for non-sinusoidal modulation in sound, although it was reduced compared to that for sinusoidal modulation (Figure 5).

In contrast, the sustained neural response was increased for non-sinusoidal compared to sinusoidal modulations in sounds, in younger adults at least. Some previous work has suggested that the sustained response increases with the degree of temporal regularity in sounds (Gutschalk et al., 2002; Herrmann and Johnsrude, 2018), and that the degree of predictability in regular repeating sounds may increase the sustained response (Heilbron and Chait, 2018). However, we have previously shown that the repetition of a regular auditory pattern can reduce the sustained response, possibly because novelty/interestingness is reduced (Herrmann et al., 2021). The invariant sinusoidal modulation at 4 Hz may have become less interesting over time, compared to the non-sinusoidal modulation which was more variable (although still somewhat regular).

### Age-related increase of neural responses to sound onset

Sound onsets elicited larger responses in the older compared to the younger group, consistent with a large body of work (Ross and Tremblay, 2009; Sörös et al., 2009; Lister et al., 2011; Alain et al., 2014; Bidelman et al., 2014; Parry et al., 2019; Irsik et al., 2021; Herrmann et al., 2022). A loss of inhibition and an increase in excitation in auditory cortex is thought to underlie hyperresponsivity to sound, and to be caused by peripheral hearing loss and aging processes (Caspary et al., 1995; Caspary et al., 2008; Takesian et al., 2012; Knipper et al., 2013; Auerbach et al., 2014; Ouellet and de Villers-Sidani, 2014; Zhao et al., 2016; Resnik and Polley, 2017; Salvi et al., 2017; Herrmann and Butler, 2021b). Indeed, our older group showed larger puretone average thresholds (although within the clinically “normal” range; https://www.iso.org/standard/42916.html) and more substantial high-frequency threshold shifts compared to younger adults, indicating that older adults had subclinical hearing impairment (Figure 1). Even mild damage to the auditory periphery leads to hyperexcitability in auditory cortex of animal models (Qiu et al., 2000; Hughes et al., 2010; Munguia et al., 2013).

Animal work demonstrates that hyperresponsivity in auditory cortex is not simply inherited from the auditory periphery, because the degree of responsivity following hearing impairment increases along the auditory pathway (Auerbach et al., 2014; Herrmann and Butler, 2021a). Hyperresponsivity is typically greatest in auditory cortex, while subcortical brain regions, such as the inferior colliculus, may not show hyperresponsivity or may even show hyporesponsivity (Auerbach et al., 2014; Chambers et al., 2016). This is consistent with recent work in humans, showing no difference in brainstem potentials between younger and older adults, whereas responses to sound in auditory cortex were larger for older compared to younger adults (Irsik et al., 2021). Moreover, although damage in the auditory periphery may be present in older adults with even minor hearing impairment, such peripheral changes do not lead to perceived loudness differences between older and younger adults that could drive age-related hyperresponsivity for sound levels used in the current study (Herrmann et al., 2019). The age-related increase in the magnitude of sound-onset responses thus was unlikely generated in the auditory periphery and transmitted along the auditory pathway, but is more likely due to cortical plasticity – mainly decreased inhibition – following sensory deprivation associated with peripheral dysfunction.

### Increased neural synchronization in older compared to younger adults

We observed increased neural synchronization in the older compared to the younger age group for both sinusoidal and non-sinusoidal modulations in sounds (Figure 5). The age-related increase appears to generalize across the different modulation types (AM, FM, AMFM), since we did not find related interactions with age group. The age-related increase in synchronization with sound modulations is consistent with the hyperresponsivity observed for sound-onset responses (Figure 3; see also Anderer et al., 1996; Sörös et al., 2009; Alain et al., 2012), suggesting a loss of inhibition in the auditory system (Caspary et al., 2008; Zhao et al., 2016; Herrmann and Butler, 2021a).

Previous studies investigating age-related differences in neural synchronization have mainly focused on amplitude modulations in sounds, and have demonstrated greater synchronization in older compared to younger listeners to narrow-band noises (Purcell et al., 2004; Goossens et al., 2016; Herrmann et al., 2019) and speech stimuli (Presacco et al., 2016b, a; Decruy et al., 2020; Broderick et al., 2021), consistent with the results reported here. Age differences in neural synchronization with frequency modulations in sounds have been less investigated, and the studies that did focus on slow frequency modulations have led to contradictory results (Boettcher et al., 2002; Henry et al., 2017). Our investigation demonstrates increased synchronization in older adults compared to younger adults for AM, FM, and AMFM sounds and that the increase is present regardless of the shape – sinusoidal and non-sinusoidal – of the modulation.

Given our data, it would be interesting to know whether the age-related increase in synchronization to FMs of narrow-band noise sounds would generalize to more naturalistic FMs, such as the low-frequency prosodic contour in speech or musical melodies. The current frequency modulation spanned ~1.5 octaves (here: 1760 Hz), which is a much wider frequency range than the range observed in natural prosody or musical melodies. Some previous work suggests that smaller frequency ranges may still lead to larger synchronization with sinusoidal frequency modulations in older adults (Boettcher et al., 2002). Investigating possible generalization of age-related differences in neural synchronization to these more naturalistic stimuli will be an important next research step.

### Reduced sustained neural response in older compared to younger adults

In the current study, frequency modulations and amplitude-frequency modulations in sounds resulted in a smaller sustained response in the older compared to the younger group, whereas no age-group difference was observed for AM-only stimuli. The age-related reduction in the sustained response to FMs and AMFMs was observed for both sinusoidal and non-sinusoidal modulations.

Age-related reductions in the sustained response have been observed previously for short sine tones (Pfefferbaum et al., 1979) and for regular patterns in sequences of short tone pips (Al Jaja et al., 2020; Herrmann et al., 2022). A previous study using AM sounds (without FM), observed a marginally significant reduction in the sustained response for older compared to younger adults (Herrmann et al., 2019). This weak, marginal group effect is consistent with the absence of an age-group effect in the current study (Figure 6). AM-only stimuli elicited smaller sustained responses (Figure 6) and less synchronization (Figure 5) compared to FM and AMFM sounds. Age-group differences to AM-only sounds may be obscured because of the lower response magnitudes (i.e., floor effect).

The difference between age groups in the magnitude of the sustained response was larger for non-sinusoidal than sinusoidal modulations. Since the magnitude of the sustained response did not differ between non-sinusoidal and sinusoidal modulations in older adults, this interaction is likely due to a greater sustained response for non-sinusoidal relative to sinusoidal modulations in younger adults. The apparent lack of sensitivity in older adults to the difference between sinusoidal and non-sinusoidal modulation further highlights how auditory-system sensitivity to low-frequency temporal structure in sounds is altered in older adulthood.

The sustained response has been suggested to reflect prediction-related processes (Heilbron and Chait, 2018), such that response magnitude increases with increasing predictability of the structure in sounds. However, in the current study we observed a larger sustained response for less predictable (non-sinusoidal) compared to more predictable (sinusoidal) temporal modulations in younger listeners (Figure 6). Moreover, the sustained response decreases when the same regular pattern is presented several times, that is, for more predictable (and less novel) patterns (Herrmann et al., 2021). Thus, it is hard to know what conclusion to draw from the reduced sustained response that we observed in older adults: we suggest that it may reflect that the auditory system of older adults treated the trial-by-trial repetition of temporal modulations as less novel compared to the auditory system of younger adults.

## Conclusions

We recorded magnetoencephalography to investigate age differences in neural responses to low-frequency modulations in amplitude, frequency, or combined amplitude and frequency. We observed hyperresponsivity to sound onset (independent of modulation types) in older compared to younger adults, consistent with a loss of inhibition in the aged auditory system. Neural synchronization with amplitude, frequency, and combined amplitude-frequency modulations were also increased for older compared to younger adults. This response potentiation generalized across sinusoidally and non-sinusoidally (less regular) modulation types. Despite greater synchronization, the sustained neural response was reduced in older compared to younger adults for frequency-modulated sounds, regardless of whether modulation was sinusoidal or non-sinusoidal. Our results thus demonstrate that the aged auditory system exhibits both hyperresponsiveness and hyporesponsiveness to sound features as reflected in different neural signatures. Sensitivity to regular temporal structure in sounds is important for tracking and comprehending speech in the presence of background noise, and our data may help explain difficulties that people in late middle age and at older ages experience with speech comprehension in listening situations when background noise is present.

## Acknowledgements

Research was supported by a Canadian Institutes of Health Research (MOP133450) grant to ISJ. BH was supported by a BrainsCAN postdoctoral fellowship (Canada First Research Excellence Fund; CFREF) and the Canada Research Chair program. We thank the Max Planck Institute for Human Cognitive and Brain Sciences for the opportunity to record the data. We thank Yvonne Wolff-Rosier for help during data acquisition.

## Author contributions

BH conceptualized and designed the study, recorded data, analyzed the data, interpreted the results, and wrote the manuscript. BM analyzed the data, interpreted the results, and edited the manuscript. ISJ conceptualized and designed the study, interpreted the results, and edited the manuscript.

## References

Al Jaja A, Grahn JA, Herrmann B, MacDonald PA (2020) The effect of aging, Parkinson’s disease, and exogenous dopamine on the neural response associated with auditory regularity processing. Neurobiology of Aging 89:71–82.

Alain C, McDonald K, Van Roon P (2012) Effects of age and background noise on processing a mistuned harmonic in an otherwise periodic complex sound. Hearing Research 283:126–135.

Alain C, Roye A, Salloum C (2014) Effects of age-related hearing loss and background noise on neuromagnetic activity from auditory cortex. Frontiers in Systems Neuroscience 8:Article 8.

Anderer P, Semlitsch HV, Saletu B (1996) Multichannel auditory event-related brain potentials: effects of normal aging on the scalp distribution of N1, P2, N2 and P300 latencies and amplitudes. Electroencephalography and clinical Neurophysiology 99:458–472.

Anderson S, Parbery-Clark A, White-Schwoch T, Kraus N (2012) Aging Affects Neural Precision of Speech Encoding. The Journal of Neuroscience 32:14156–14164.

Auerbach BD, Rodrigues PV, Salvi RJ (2014) Central gain control in tinnitus and hyperacusis. Frontiers in Neurology 5:Article 206.

Barascud N, Pearce MT, Griffiths TD, Friston KJ, Chait M (2016) Brain responses in humans reveal ideal observer-like sensitivity to complex acoustic patterns. Proceedings of the National Academy of Sciences 113:E616–E625.

Barnes R, Jones MR (2000) Expectancy, Attention, and Time. Cognitive Psychology 41:254–311.

Bauer A-KR, van Ede F, Quinn AJ, Nobre AC (2021) Rhythmic Modulation of Visual Perception by Continuous Rhythmic Auditory Stimulation. The Journal of Neuroscience 41:7065–7075.

Bell AJ, Sejnowski TJ (1995) An information maximization approach to blind separation and blind deconvolution. Neural Computation 7:1129–1159.

Bendixen A (2014) Predictability effects in auditory scene analysis: a review. Frontiers in Neuroscience 8:Article 60.

Bidelman GM, Villafuerte JW, Moreno S, Alain C (2014) Age-related changes in the subcorticalecortical encoding and categorical perception of speech. Neurobiology of Aging 35:2526–2540.

Boettcher FA, Madhotra D, Poth EA, Mills JH (2002) The frequency-modulation following response in young and aged human subjects. Hearing Research 165:10–18.

Brodbeck C, Simon JZ (2020) Continuous speech processing. Current Opinion in Physiology 18:25–31.

Broderick MP, Di Liberto GM, Anderson AJ, Rofes A, Lalor EC (2021) Dissociable electrophysiological measures of natural language processing reveal differences in speech comprehension strategy in healthy ageing. Scientific Reports 11:4963.

Caspary DM, Milbrandt JC, Helfert RH (1995) Central auditory aging: GABA changes in the inferior colliculus. Experimental Gerontology 30:349–360.

Caspary DM, Ling L, Turner JG, Hughes LF (2008) Inhibitory neurotransmission, plasticity and aging in the mammalian central auditory system. The Journal of Experimental Biology 211:1781–1791.

Chambers AR, Salazar JJ, Polley DB (2016) Persistent Thalamic Sound Processing Despite Profound Cochlear Denervation. Frontiers in Neural Circuits 10:Article 72.

Cruickshanks KJ, Wiley TL, Tweed TS, Klein BEK, Klein R, Mares-Perlman JA, Nondahl DM (1998) Prevalence of Hearing Loss in Older Adults in Beaver Dam, Wisconsin. American Journal of Epidemiology 148:879–886.

Decruy L, Vanthornhout J, Francart T (2020) Hearing impairment is associated with enhanced neural tracking of the speech envelope. Hearing Research 393:107961.

Dheerendra P, Barascud N, Kumar S, Overath T, Griffiths TD (2021) Dynamics underlying auditory-object-boundary detection in primary auditory cortex. European Journal of Neuroscience 54:7274–7288.

Edwards E, Chang EF (2013) Syllabic (~2-5 Hz) and fluctuation (~1-10 Hz) ranges in speech and auditory processing. Hearing Research 305:113–134.

Fastl H (1982) Fluctuation strength and temporal masking patternsof amplitude-modulated broadband noise. Hearing Research 8:59–69.

Fastl H (1983) Fluctuation Strength of Modulated Tones and Broadband Noise. In: HEARING — Physiological Bases and Psychophysics (Klinke R, Hartmann R, eds), pp 282–288. Berlin, Heidelberg: Springer Berlin Heidelberg.

Fastl H, Zwicker E (2007) Fluctuation Strength. In: Psychoacoustics: Facts and Models (Fastl H, Zwicker E, eds), pp 247-256. Berlin, Heidelberg: Springer Berlin Heidelberg.

Feder K, Michaud D, Ramage-Morin P, McNamee J, Beauregard Y (2015) Prevalence of hearing loss among Canadians aged 20 to 79: Audiometric results from the 2012/2013 Canadian Health Measures Survey. Health Reports 26:18–25.

Fuglsang SA, Märcher-Rørsted J, Dau T, Hjortkjær J (2020) Effects of sensorineural hearing loss on cortical synchronization to competing speech during selective attention. The Journal of Neuroscience:1936–1919.

Giraud A-L, Poeppel D (2012) Cortical oscillations and speech processing: emerging computational principles and operations. Nature Neuroscience 15.

Goman AM, Lin FR (2016) Prevalence of Hearing Loss by Severity in the United States. American Journal of Public Health 106:1820–1822.

Goossens T, Vercammen C, Wouters J, van Wieringen A (2016) Aging Affects Neural Synchronization to Speech-Related Acoustic Modulations. Frontiers in Aging Neuroscience 8:Article 133.

Goossens T, Vercammen C, Wouters J, Van Wieringen A (2019) The association between hearing impairment and neural envelope encoding at different ages. Neurobiology of Aging 74:202–212.

Greenhouse SW, Geisser S (1959) On Methods in the Analysis of Profile Data. Psychometrika 24:95–112.

Gutschalk A, Uppenkamp S (2011) Sustained responses for pitch and vowels map to similar sites in human auditory cortex. NeuroImage 56:1578–1587.

Gutschalk A, Patterson RD, Rupp A, Uppenkamp S, Scherg M (2002) Sustained Magnetic Fields Reveal Separate Sites for Sound Level and Temporal Regularity in Human Auditory Cortex. NeuroImage 15:207–216.

Hämäläinen MS, Hari R (2002) Magnetoencephalographic (MEG) Characterization of Dynamic Brain Activation: Basic Principles and Methods of Data Collection and Source Analysis. In: Brain Mapping: The Methods (Toga AW, Mazziotta JC, eds), pp 227–253: Academic Press.

Hämäläinen MS, Hari R, Ilmoniemi RJ, Knuutila J, Lounasmaa OV (1993) Magnetoencephalography – theory, instrumentation, and applications to noninvasive studies of the working human brain. Reviews of Modern Physics 65:413–497.

Harris KC, Wilson S, Eckert MA, Dubno JR (2012) Human evoked cortical activity to silent gaps in noise: effects of age, attention, and cortical processing speed. Ear Hear 33:330–339.

Heilbron M, Chait M (2018) Great expectations: Is there evidence for predictive coding in auditory cortex? Neuroscience 389:54–73.

Henry MJ, Obleser J (2012) Frequency modulation entrains slow neural oscillations and optimizes human listening behavior. Proceedings of the National Academy of Sciences 109:20095–20100.

Henry MJ, Herrmann B (2014) Low-Frequency Neural Oscillations Support Dynamic Attending in Temporal Context. Timing & Time Perception 2:62–86.

Henry MJ, Herrmann B, Obleser J (2014) Entrained neural oscillations in multiple frequency bands co-modulate behavior. Proceedings of the National Academy of Sciences 111:14935–14940.

Henry MJ, Herrmann B, Kunke D, Obleser J (2017) Aging affects the balance of neural entrainment and top-down neural modulation in the listening brain. Nature Communications 8:15801.

Herrmann B, Johnsrude IS (2018) Neural signatures of the processing of temporal patterns in sound. The Journal of Neuroscience 38:5466–5477.

Herrmann B, Butler BE (2021a) Hearing Loss and Brain Plasticity: The Hyperactivity Phenomenon. Brain Structure & Function 226:2019–2039.

Herrmann B, Butler BE (2021b) Aging auditory cortex: The impact of reduced inhibition on function. In: Assessments, Treatments and Modelling in Aging and Neurological Disease: The Neuroscience of Aging (Martin CR, Preedy VR, Rajendram R, eds), pp 183–192: Academic Press.

Herrmann B, Maess B, Johnsrude IS (2018) Aging Affects Adaptation to Sound-Level Statistics in Human Auditory Cortex. The Journal of Neuroscience 38:1989–1999.

Herrmann B, Buckland C, Johnsrude IS (2019) Neural signatures of temporal regularity processing in sounds differ between younger and older adults. Neurobiology of Aging 83:73–85.

Herrmann B, Araz K, Johnsrude IS (2021) Sustained neural activity correlates with rapid perceptual learning of auditory patterns. NeuroImage 238:118238.

Herrmann B, Maess B, Johnsrude IS (2022) A Neural Signature of Regularity in Sound is Reduced in Older Adults. Neurobiology of Aging 109:1–10.

Herrmann B, Henry MJ, Grigutsch M, Obleser J (2013) Oscillatory Phase Dynamics in Neural Entrainment Underpin Illusory Percepts of Time. The Journal of Neuroscience 33:15799–15809.

Hughes LF, Turner JG, Parrish JL, Caspary DM (2010) Processing of broadband stimuli across A1 layers in young and aged rats. Hearing Research 264:79–85.

Irsik VC, Almanaseer A, Johnsrude IS, Herrmann B (2021) Cortical Responses to the Amplitude Envelopes of Sounds Change with Age. The Journal of Neuroscience 41:5045–5055.

John MS, Dimitrijevic A, van Roon P, Picton TW (2001) Multiple Auditory Steady-State Responses to AM and FM Stimuli. Audiology & Neuro-Otology 6:12–27.

Jones MR, Boltz MG (1989) Dynamic Attending and Responses to Time. Psychological Review 96:459–491.

Jones MR, Moynihan H, MacKenzie N, Puente J (2002) Temporal Aspects of Stimulus-Driven Attending in Dynamic Arrays. Psychological Science 13:313–319.

Keitel C, Thut G, Gross J (2016) Visual cortex responses reflect temporal structure of continuous quasi-rhythmic sensory stimulation. NeuroImage 146:58–70.

Knipper M, Van Dijk P, Nunes I, Rüttiger L, Zimmermann U (2013) Advances in the neurobiology of hearing disorders: Recent developments regarding the basis of tinnitus and hyperacusis. Progress in Neurobiology 111:17–33.

Lachaux J-P, Rodriguez E, Martinerie J, Varela FJ (1999) Measuring Phase Synchrony in Brain Signals. Human Brain Mapping 8:194–208.

Lakatos P, Karmos G, Mehta AD, Ulbert I, Schroeder CE (2008) Entrainment of Neuronal Oscillations as a Mechanism of Attentional Selection. Science 320:110–113.

Lakatos P, Musacchia G, O’Connel MN, Falchier AY, Javitt DC, Schroeder CE (2013) The Spectrotemporal Filter Mechanism of Auditory Selective Attention. Neuron 77:750–761.

Lister JJ, Maxfield ND, Pitt GJ, Gonzalez VB (2011) Auditory evoked response to gaps in noise: older adults. International Journal of Audiology 50:211–225.

Maiste A, Picton TW (1989) Human auditory evoked potentials to frequency-modulated tones. Ear & Hearing 10:153–160.

Makeig S, Bell AJ, Jung T-P, Sejnowski TJ (1996) Independent component analysis of electroencephalographic data. In: Advances in Neural Information Processing Systems (Touretzky D, Mozer M, Hasselmo M, eds). Cambridge, MA, USA: MIT Press.

Moore BCJ (2007) Cochlear Hearing Loss: Physiological, Psychological and Technical Issues. West Sussex, Engand: John Wiley & Sons, Ltd.

Moore BCJ, Wojtczak M, Vickers D (1996) Effect of loudness recruitment on the perception of amplitude modulation. Journal of the Acoustical Society of America 100:481–489.

Munguia R, Pienkowski M, Eggermont JJ (2013) Spontaneous firing rate changes in cat primary auditory cortex following long-term exposure to non-traumatic noise: Tinnitus without hearing loss? Neuroscience Letters 546:46–50.

Nobre AC, van Ede F (2018) Anticipated moments: temporal structure in attention. Nature Reviews Neuroscience 19:34–48.

Nolte G, Bai O, Wheaton L (2004) Identifying true brain interaction from EEG data using the imaginary part of coherency. Clinical Neurophysiology 115:2292–2307.

Nozaradan S, Peretz I, Missal M, Mouraux A (2011) Tagging the Neuronal Entrainment to Beat and Meter. The Journal of Neuroscience 31:10234–10240.

O’Connell MN, Barczak A, Schroeder CE, Lakatos P (2014) Layer Specific Sharpening of Frequency Tuning by Selective Attention in Primary Auditory Cortex. The Journal of Neuroscience 34:16496–16508.

Obleser J, Kayser C (2019) Neural Entrainment and Attentional Selection in the Listening Brain. Trends in Cognitive Sciences 23:913–926.

Obleser J, Herrmann B, Henry MJ (2012) Neural oscillations in speech: don’t be enslaved by the envelope. Frontiers in Human Neuroscience 6:Article 250.

Oostenveld R, Fries P, Maris E, Schoffelen JM (2011) FieldTrip: Open source software for advanced analysis of MEG, EEG, and invasive electrophysiological data. Computational Intelligence and Neuroscience 2011:Article ID 156869.

Ouellet L, de Villers-Sidani E (2014) Trajectory of the main GABAergic interneuron populations from early development to old age in the rat primary auditory cortex. Frontiers in Neuroanatomy 8:Article 40.

Parry LV, Maslin MRD, Schaette R, Moore DR, Munro KJ (2019) Increased auditory cortex neural response amplitude in adults with chronic unilateral conductive hearing impairment. Hearing Research 372:10–16.

Peelle JE, Davis MH (2013) Neural oscillations carry speech rhythm through to comprehension. Frontiers in Psychology 3:Article 320.

Peelle JE, Gross J, Davis MH (2013) Phase-Locked Responses to Speech in Human Auditory Cortex are Enhanced During Comprehension. Cerebral Cortex 23:1378–1387.

Pfefferbaum A, Ford JM, Roth WT, Hopkins WF, Kopell BS (1979) Event-related potential changes in healthy aged females. Electroencephalography and Clinical Neurophysiology 46:81–86.

Pichora-Fuller MK (2003) Processing speed and timing in aging adults: psychoacoustics, speech perception, and comprehension. International Journal of Audiology 42:S59–S67.

Pichora-Fuller MK, Kramer SE, Eckert MA, Edwards B, Hornsby BWY, Humes LE, Lemke U, Lunner T, Matthen M, Mackersie CL, Naylor G, Phillips NA, Richter M, Rudner M, Sommers MS, Tremblay KL, Wingfield A (2016) Hearing Impairment and Cognitive Energy: The Framework for Understanding Effortful Listening (FUEL). Ear & Hearing 37 Suppl 1:5S–27S.

Picton TW, Woods DL, Proulx GB (1978) Human auditory sustained potentials. II. Stimulus relationships. Electroencephalography and clinical Neurophysiology 45:198–210.

Picton TW, Hillyard SA, Krausz HI, Galambos R (1974) Human auditory evoked potentials. I: Evaluation of components. Electroencephalography and clinical Neurophysiology 36:179–190.

Picton TW, John SM, Dimitrijevic A, Purcell DW (2003) Human auditory steady-state responses. International Journal of Audiology 42:177–219.

Plack CJ (2014) The sense of hearing. New York, USA: Psychology Press.

Polich J, Aung M, Dalessio DJ (1988) Long Latency Auditory Evoked Potentials: Intensity, Inter-Stimulus Interval, and Habituation. The Pavlovian Journal of Biological Science 23:35–40.

Presacco A, Simon JZ, Anderson S (2016a) Effect of informational content of noise on speech representation in the aging midbrain and cortex. Journal of Neurophysiology 116:2356–2367.

Presacco A, Simon JZ, Anderson S (2016b) Evidence of degraded representation of speech in noise, in the aging midbrain and cortex. Journal of Neurophysiology 116:2346–2355.

Presacco A, Simon JZ, Anderson S (2019) Speech-in-noise representation in the aging midbrain and cortex: Effects of hearing loss. PLoS ONE 14:e0213899.

Purcell DW, John SM, Schneider BA, Picton TW (2004) Human temporal auditory acuity as assessed by envelope following responses. The Journal of the Acoustical Society of America 116:3581–3593.

Qiu C, Salvi R, Ding D, Burkard R (2000) Inner hair cell loss leads to enhanced response amplitudes in auditory cortex of unanesthetized chinchillas: evidence for increased system gain. Hearing research 139:153–171.

Resnik J, Polley DB (2017) Fast-spiking GABA circuit dynamics in the auditory cortex predict recovery of sensory processing following peripheral nerve damage. eLife 6:e21452.

Rosen S (1992) Temporal Information in Speech: Acoustic, Auditory and Linguistic Aspects. Philosophical Transactions: Biological Sciences 336:367–373.

Ross B, Tremblay KL (2009) Stimulus experience modifies auditory neuromagnetic responses in young and older listeners. Hearing Research 248:48–59.

Ross B, Picton TW, Pantev C (2002) Temporal integration in the human auditory cortex as represented by the development of the steady-state magnetic field. Hearing Research 165:68–84.

Ross B, Schneider B, Snyder JS, Alain C (2010) Biological Markers of Auditory Gap Detection in Young, Middle-Aged, and Older Adults. PLOS ONE 5:e10101.

Salvi R, Sun W, Ding D, Chen G-D, Lobarinas E, Wang J, Radziwon K, Auerbach BD (2017) Inner Hair Cell Loss Disrupts Hearing and Cochlear Function Leading to Sensory Deprivation and Enhanced Central Auditory Gain. Frontiers in Neuroscience 10:Article 621.

Schadow J, Lenz D, Thaerig S, Busch NA, Fründ I, Herrmann CS (2007) Stimulus intensity affects early sensory processing: Sound intensity modulates auditory evoked gamma-band activity in human EEG. International Journal of Psychophysiology 65:152–161.

Schröger E (2005) The Mismatch Negativity as a Tool to Study Auditory Processing. Acta Acustica united with Acustica 91:490–501.

Schröger E (2007) Mismatch Negativity: A Microphone into Auditory Memory. Journal of Psychophysiology 21:138–146.

Snyder JS, Alain C (2007) Toward a neurophysiological theory of auditory stream segregation. Psychological Bulletin 133:780–799.

Sohoglu E, Chait M (2016) Detecting and representing predictable structure during auditory scene analysis. eLife 5:e19113.

Sörös P, Treismann IK, Manemann E, Lütkenhöner B (2009) Auditory temporal processing in healthy aging: a magnetoencephalographic study. BMC Neuroscience 10:34.

Southwell R, Chait M (2018) Enhanced deviant responses in patterned relative to random sound sequences. Cortex 109:92–103.

Southwell R, Baumann A, Gal C, Barascud N, Friston KJ, Chait M (2017) Is predictability salient? A study of attentional capture by auditory patterns. Philosophical Transactions of the Royal Society B 372:20160105.

Stefanics G, Hangya B, Hernádi I, Winkler I, Lakatos P, Ulbert I (2010) Phase Entrainment of Human Delta Oscillations Can Mediate the Effects of Expectation on Reaction Speed. The Journal of Neuroscience 30:13578–13585.

Takesian AE, Kotak VC, Sanes DH (2012) Age-dependent effect of hearing loss on cortical inhibitory synapse function. Journal of Neurophysiology 107:937–947.

Taulu S, Kajola M, Simola J (2004) Suppression of Interference and Artifacts by the Signal Space Separation Method. Brain Topography 16:269–275.

Taulu S, Simola J, Kajola M (2005) Applications of the Signal Space Separation Method. IEEE Transactions On Signal Processing 53:3359–3372.

Teki S, Barascud N, Picard S, Payne C, Griffiths TD, Chait M (2016) Neural Correlates of Auditory Figure-Ground Segregation Based on Temporal Coherence. Cerebral Cortex 26:3669–3680.

Varnet L, Ortiz-Barajas CM, Erra RG, Gervain J, Lorenzi C (2017) A cross-linguistic study of speech modulation spectra. The Journal of the Acoustical Society of America 142:1976–1989.

Vecchi AO, León RG, Kohlrausch A (2016) Modelling the sensation of fluctuation strength. Proceedings of Meetings on Acoustics 28:050005.

Winkler I, Denham SL, Nelken I (2009) Modeling the auditory scene: predictive regularity representations and perceptual objects. Trends in Cognitive Sciences 13:532–540.

Zhao Y, Song Q, Li X, Li C (2016) Neural Hyperactivity of the Central Auditory System in Response to Peripheral Damage. Neural Plasticity 2016:2162105.

